# Autophagy promotes tumor growth through facilitating JAK/STAT signaling in a lysosomal degradation independent manner

**DOI:** 10.64898/2026.02.12.705500

**Authors:** András Rubics, Natali Neuhauser, Dorottya Károlyi, Bálint Sólyom Bótor, Fergal O’Farrell, Szabolcs Takáts

## Abstract

Autophagy contributes to normal cells’ physiology and is essential for progression of malignant tumors. While autophagy is mostly considered as a self-degradative and self-renewal process, it has non-degradative functions whose contribution to tumor progression is poorly explored. Here we use the autophagy dependent Drosophila Ras^V12^, Scrib^-/-^ carcinoma model to examine whether perturbation of distinct steps of autophagy differentially influences tumor progression. We found that inhibition of autophagosome formation, by mutating Atg13 or Atg6 either in the tumor or in the whole animal significantly decreased tumor growth. In contrast, blocking the later autophagosome-lysosome fusion (by loss of Vps39 or Syx17) and thereby autolysosomal degradation, does not reduce tumor size. We observed that an early (Atg13), but not a late (Vps39 or Syx17) block in autophagy showed reduced activity of JAK/STAT signaling, known to be critical for the progression of this tumor type. Importantly, we demonstrated that both Atg13 and Vps39 deficient tumors accumulated Stat92E inhibitor Su(var)2-10/dPIAS, a recently identified autophagic cargo, however in Vps39 mutants Su(var)2-10 is sequestered into autophagosomes. Finally, we found that reduction of Su(var)2-10 partially restores JAK/STAT signaling and rescues the growth of Atg13-deficient tumors, indicating its sequestration is a crucial mechanism to promote tumor progression.

## Introduction

The emergence of malignant cancer cells is accompanied by reprogramming the metabolism of both host and tumor tissue. As a part of tumor associated metabolic alterations multiple tumors become highly dependent on macroautophagy (hereafter referred as autophagy), during which portions of cytoplasmic material are degraded in lysosomes, and the resulting building blocks can be reused by the cell^1^. The integrity of autophagic pathway has great importance in the maintenance of proper cellular functions, while dysregulation of autophagy contributes to various pathologies^2^. Cancer cells defective in autophagy are not able to grow properly ^3-5^ and autophagy inhibition can be a potent antitumor strategy^1^. Interestingly, tumors can also become dependent on autophagic activity of the host tissues and earlier studies showed that loss of autophagy in the host tissues can suppress tumor growth and cancer cachexia ^5-7^.

The importance of autophagy for tumor progression is still incompletely understood. The most generally approved explanation is that cancer cells require large amount of nutrients to support their anabolic processes, and autophagic degradation of superfluous organelles in the cancer cell itself, or in host tissues, can provide organic monomers that can be consumed by the tumors^1,5,7-9^. However, in addition to releasing nutrients, autophagy has other less well characterized functions too. Autophagy can regulate several signaling or metabolic processes, by sequestering and hence isolating signaling proteins from the cytosol^10-12^, by enwrapping them into autophagosomes, and eventually degrading them in autolysosomes.

Autophagy is a complex multi-step pathway. It begins with the formation of a cistern-like, double membrane structure, called phagophore that sequesters cytoplasmic material or even complete organelles and after sealing the phagophore’s edges the cargo gets enwrapped into a double-membraned autophagosome. These transport vesicles then fuse with lysosomes, where the cargo gets degraded and the released building blocks can be reused. Each progressive step in this process can be blocked via genetic means, classically used to study autophagy, less so for tumor progression. Early steps of autophagy, the initiation and completion of the phagophore and the autophagosome formation is regulated by four highly conserved complexes formed by Autophagy-related (Atg) genes^1^. Then, the maturation of the newly formed autophagosome and its subsequent fusion with lysosomes is mediated by the HOPS tethering complex (composed of Vps11, Vps16A, Vps18, Vps33A, Vps39 and Vps41 proteins), a SNARE complex (formed by Syntaxin 17, Snap29, Vamp7/8 SNARE proteins) and several small GTPases (Rab2, Rab7, Arl8)^13^. Autophagy is mostly considered as a degradative process and each of these regulatory steps are required for efficient cargo degradation. However, it is still poorly understood whether impairment of distinct regulatory steps of autophagy (like autophagosome formation or autophagosome-lysosome fusion) are equally important for cancer progression.

Studies with Ras^V12^; Scrib^-/-^ *Drosophila* carcinoma model induced in the larval developing eye pointed out that tumor specific loss of autophagic genes, like Atg1, Atg13 can critically decrease the growth of malignant carcinoma^4,14^. Moreover, extensive levels of autophagy are induced in host tissues in the presence of the same or similar tumors^5,6,9^ and the lack of Atg13 or Atg14 in the entire body (systemic loss of autophagy) still results in reduced tumor growth^5,6^. However, the role of autophagosome-lysosome fusion in the progression of this tumor type has not been tested. Hence, we use the same *Drosophila* tumor model combined with mutations of distinct steps of the autophagic process. This approach enabled us to elucidate whether the suppression of tumor growth observed in autophagosome formation defective Atg mutants is really due to impaired autophagic degradation and nutrient release, and to genetically dissect the respective contributions of the autophagosome formation and autophagosome-lysosome fusion steps to tumor progression.

## Results

### Only early steps of autophagy are critical for tumor progression

To assay whether cancer cell specific autophagy is contributing to the progression of autophagy dependent tumors as a degradative process we used the well-established *Drosophila* carcinoma model, where malignant cell clones, which simultaneously overexpress the *Ras*^*V12*^ oncogene and are homozygous mutant for the *scrib* cell polarity gene, are induced by MARCM strategy, that eventually develop to an aggressive carcinoma^15^ (hereafter we refer to these as Ras^V12^, Scrib^-/-^/RS tumors). We chose to test the effect of the tumor specific loss of genes that are well known regulators of distinct steps of autophagy. Atg13 and Atg6 proteins are members of Atg1 (homolog of ULK1/2) and Vps34 kinase complexes respectively and both act as critical upstream regulators of phagophore and autophagosome formation. Importantly, it was demonstrated before that cell autonomous loss of Atg6 and Atg13 genes results in the complete lack of autophagosomes^16,17^. In contrast, Vps39 is a subunit of the HOPS tethering complex, an essential regulator of autophagosome lysosome fusion, and the lack of this gene results in the accumulation of autophagosomes that are unable to fuse with lysosomes^18^. Importantly, loss of any of these genes prevents the formation of autolysosomes and autophagic degradation. In line with other previous findings^4,5^, generating RS tumors that were also homozygous for a mutant allele of Atg13^16^ or Atg6^17^ in otherwise heterozygous larvae resulted in tumors with extremely reduced size compared to the same aged control RS ones **(Figure 1A, B, D, Supplementary Figure S1A-C)**. Conversely, cancer cell specific loss of Vps39 did not affect the progression of RS tumors **(Figure 1C, D)**. To verify that the Vps39 mutant allele^19^ we are using really inhibits autophagic degradation we carried out an autophagic flux assay^6^, that measures the lysosomal hydrolase mediated cleavage of mCherry-GFP-Atg8a reporter. Western blotting of lysates of tumors which expressed mCherry-GFP-Atg8a in a cancer cell specific manner revealed that RS tumors showed high levels of free mCherry indicating that autophagic degradation is active in them **(Figure 1E)**. In contrast, free mCherry was almost undetectable in Atg13, Atg6 or Vps39 mutant tumor samples **(Figure 1E, Supplementary Figure S1D)** proving that these tumors are defective for autophagic degradation. As further verification that autophagosome formation and fusion is blocked in Atg13 and Vps39 mutant tumors respectively, we tested the subcellular distribution of phagophores and autophagosomes by immunolabeling autophagic membrane marker Atg8a (*Drosophila* ortholog of LC3). In control tumors Atg8a localized to puncta-like structures **(Figure 1F, I)** indicating the presence of a moderate amount of autophagic vesicles in this tissue. As expected, the level of punctate autophagic structures strongly decreased, while the cytoplasmic background of Atg8a was visibly increased in Atg13 mutants **(Figure 1G, I)**, demonstrating the impairment to autophagosome formation in this genotype. On the other hand, Vps39 mutants showed a massive accumulation of puncta-like Atg8a positive autophagosomes **(Figure 1H, I)**, a phenotype that observed upon autophagosome-lysosome fusion deficient conditions^18^. Taken together with the tumor growth data, these findings raise the possibility that cancer cells require only early steps of autophagy, but not autophagic degradation, to maintain their progression.

**Figure 1.**
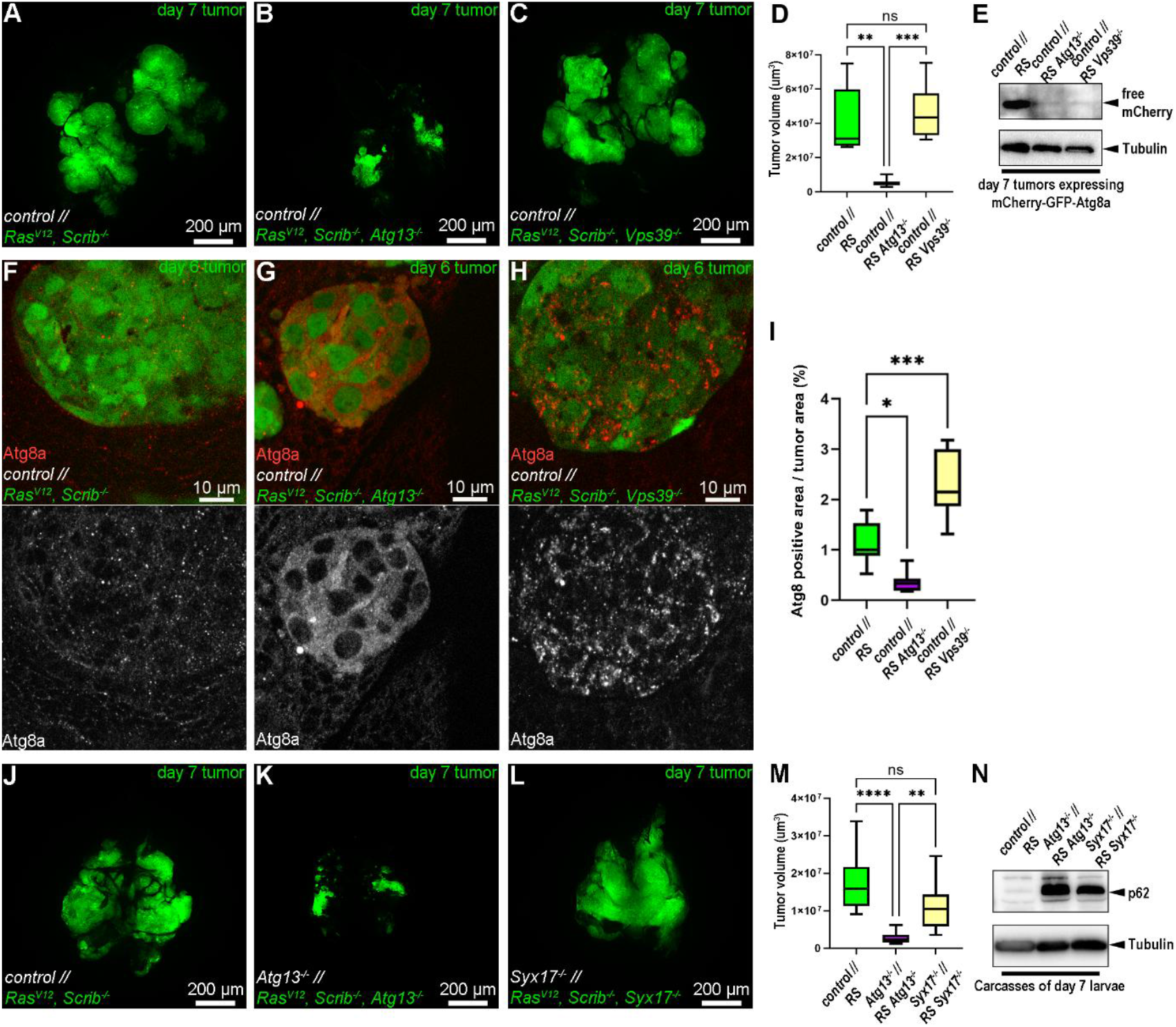
Autophagosome formation but not autophagsome-lysosome fusion is required for the growth of RS tumors. **(A-C)** Representative images about control **(A)**, Atg13^-/-^ **(B)** and Vps39^-/-^ **(C)** RS tumors that were induced in larvae with control background. **(D)** Quantification of data presented in A-C. 7-9 tumors/genotype were analyzed, n=7 (A), 9 (B), 8 (C). Kruskal-Wallis tests,**: p<0,01, ***: p<0,001. **(E)** Western blotting of day 7 tumors monitoring autophagy viaprocessed free mCherry, the lysosomal cleavage product of mCherry-GFP-Atg8a reporter, in Atg13 and Vps39 deficient tumors. **(F-H)** Anti-Atg8a immunostaining on day 6 control **(F)**, Atg13**(G)** and Vps39 **(H)** mutant RS tumors that were induced in larvae with control genetic background. **(I)** Quantification of data presented on F-H. 7-8 tumors/genotype were analyzed, n=8 (F), 7 (G), 8 (H). one-way ANOVA, *: p<0,05, ***: p<0,001. **(J-L)** Representative images ofcontrol **(J)**, Atg13^-/-^ **(K)** and Syx17^-/-^ **(L)** RS tumors that were induced in larvae in either control **(J)**, Atg13^-/-^ **(K)** or Syx17^-/-^ **(L)** backgrounds respectively. **(M)** Quantification of data presented in J-L. 13-18 tumors/genotype were analyzed, n=18 (J), 13 (K), 15 (L). Kruskal-Wallis tests, ns: notsignificant, **: p<0,01, ****: p<0,0001. **(N)** Western blot detection of p62 and Atg8a autophagy substrates (lower levels indicate higher autophagy) in the lysates of body wall muscles from Atg13 and Syx17 mutant or control day 7 larvae that carry RS tumors. Scales bars as indicated.

It was previously demonstrated that the progression of RS tumors is dependent on the autophagic activity of both the tumor itself and the non-transformed host tissues like microenvironment, adipose tissue or body wall muscles^5,6^. To characterize whether autophagy in host tissues is contributing to tumor progression via a degradative process we generated tumors in larvae whose entire body (including tumor and host tissues) was defective for autophagosome formation (Atg13^-/-^) or autophagosome-lysosome fusion (Syntaxin17/Syx17^-/-^)^20^. As expected, tumors raised in Atg13 mutant larvae did not progress like control RS tumors **(Figure 1J, K, M)**, while tumor growth in a Syx17 mutant background progressed similarly to RS controls **(Figure 1L, M)**. Moreover, western blotting showed that muscles from Atg13 and Syx17 mutant larvae showed accumulation of the specific autophagic cargo p62 **(Figure 1N)** indicating that loss of either of these genes strongly perturbs autophagic degradation in host tissues. This finding suggests autophagy has an alternative function both in the tumor and host tissues, that requires early steps of autophagosome formation, but not lysosomal fusion and cargo degradation.

### Autophagosome formation is critical for cancer cell division and survival

The difference in progression capability between Atg13 and Vps39 mutant tumors clearly pointed out that autophagosome formation and autophagosome-lysosome fusion are differentially contributing to tumor cell physiology. To uncover why autophagosome formation deficient tumors cannot progress we analyzed cancer cell proliferation and viability by immunolabeling of dividing cells by Phospho-Histone H3 (P-H3) and apoptotic ones by with anti-Cleaved Death Caspase-1 (Dcp-1). This experiment revealed that tumor specific loss of Atg13 but not Vps39 significantly decreases the proliferation of cancer cells compared to control tumor tissues **(Figure 2A-D)**. Additionally, compared to cancer cells of control RS tumors **(Fig. 2E, H)** where large apoptotic zones were mostly visible in the microenvironment^21^ in a close vicinity to the border of tumor/microenvironment tissue areas, Atg13 mutant tumor cell clones **(Fig 2F, H)** showed remarkable overlap with apoptotic marker Dcp-1. In contrast, the apoptotic ratio remained moderate in Vps39 mutant malignant tissues **(Figure 2G, H)** and showed similar pattern to control RS ones **(Figure 2E, G)**. These findings point out that early steps of autophagy, but not autophagosome-lysosome fusion and subsequent degradation, are critical for tumor cell proliferation and survival.

**Figure 2.**
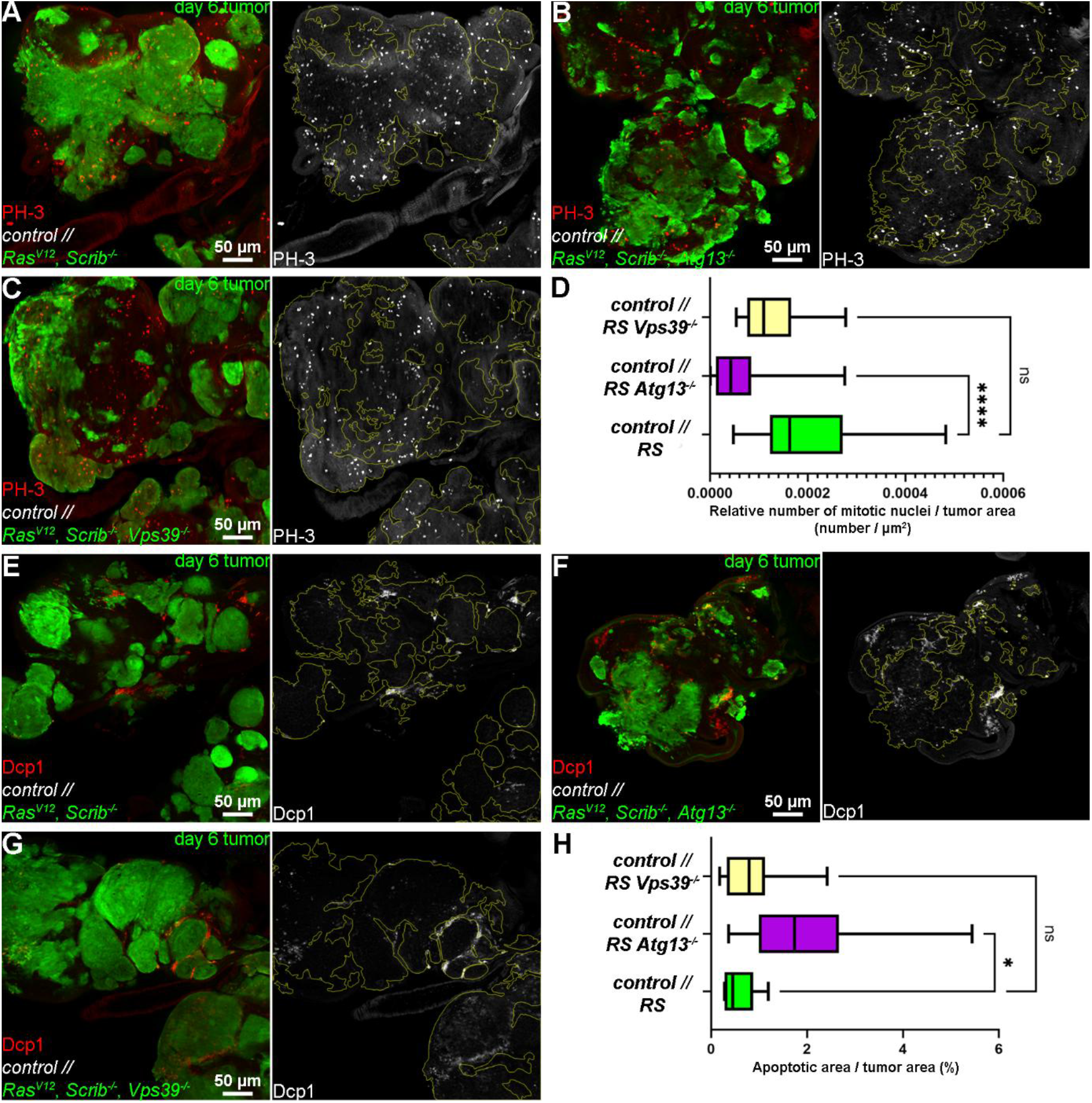
Autophagosome formation but not autophagosome-lysosome fusion is required for the proliferation of RS cancer cells. **(A-C)** Immunolabeling with anti-Phospho-Histone H3 (P-H3) of day 7 eye discs to detect levels of mitosis in control **(A)**, Atg13^-/-^ **(B)** or Vps39^-/-^ **(C)** mutant tumors. Yellow bounding box marks borders between GFP+ tumor tissue and GFP negative control tissue areas. **(D)** Quantification of data presented in A-C where mitosis from specifically GFP+ tissue were compared. 12-18 eye discs/genotype were analyzed, n=18 (A), 16(B), 12 (C). Kruskal-Wallis tests, ns: not significant, ****: p<0,0001. **(E-G)** Immunolabeling of eye discs with cleaved form of effector caspase Dcp-1 to detect levels of apoptosis in control RS **(E)** or Atg13^-/-^**(F)** and Vps39^-/-^ **(G)** tumors. **(H)** Quantification of data presented in E-G. Intensity of labelling and hence levels of apoptosis within tumor specific GFP+ tissue (yellow bounding box of genotypes indicated) were analyzed. 5-12 eye discs/genotype were analyzed, n=5 (E), 10 (F), 12 (G). Kruskal-Wallis tests, ns: not significant.

### Early steps of autophagy are required for JAK/STAT signaling

The requirement of Atg13 but not Vps39 for tumor cell proliferation and evading apoptosis raised the possibility that early steps of autophagy might be critical for the activity of a pro-tumorigenic signaling pathway. It was described previously that the progression of RS tumors is highly dependent on the activation of JAK/STAT signaling^14,15,22^. Moreover, the contribution of autophagy for upregulation of JAK/STAT was also reported in several human cancer types^23-25^. To test whether manipulation of autophagy is affecting JAK/STAT signaling, we generated RFP labelled control and autophagy deficient tumors and analyzed the activity of JAK/STAT, by using a GFP reporter whose expression is specifically driven by the activated Stat92E transcription factor^26^. As expected, eye discs carrying RS tumors showed high Stat92E driven GFP signal, while discs with Atg13 deficient tumors showed a moderate Stat92E activity **(Figure 3A, B)**. Importantly, we observed high Stat92E induced GFP signal in Vps39 mutant RS tumors **(Figure 3C)**, indicating that autolysosomal degradation is not required for JAK/STAT signaling. This finding was further confirmed by western blot analysis, which clearly showed that the lysates of eye discs that carry Atg13 and Atg6 deficient tumors express remarkably lower levels of Stat92E driven GFP compared to samples with control and Vps39 mutant tumors **(Figure 3D, Supplementary Figure S1E)**. To understand if the difference in size between Atg13 and Vps39 mutant tumors could be due to decreased JAK/STAT signaling, we tested whether early or late steps of autophagy are showing different sensitivity to the loss of the downstream JAK/STAT effector Stat92E. Importantly, tumor specific silencing of Stat92E significantly decreased the progression of both control and Vps39 mutant tumors, showing it is required for optimal tumor growth, while the small size of Atg13 deficient tumors was not further affected by the depletion of STAT signaling **(Figure 3E-M)**.

**Figure 3.**
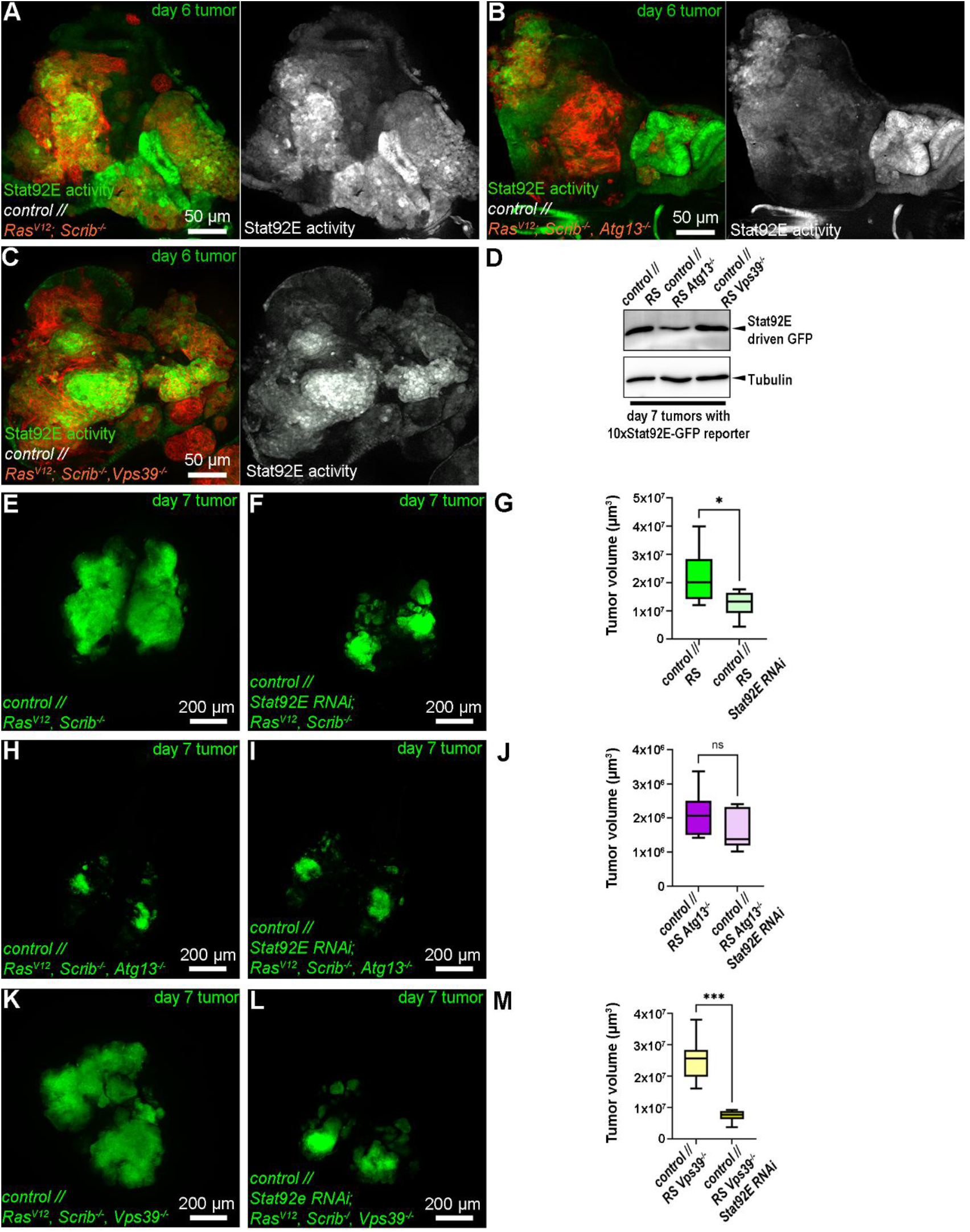
JAK/STAT signaling activity is dependent on the early steps of autophagy but not autophagosome-lysosome fusion. **(A-C)** Fluorescent Stat92E activity reporter, 10xGFP-STAT92E indicates the levels of Stat signaling in control **(A)**, Atg13 mutant **(B)** and Vps39-deficient **(C)** RS tumors. **(D)** Western blotting to detect a stat92E activity-readout from lysates of day 7 tumors of the genotypes in A-C. **(E, F)** Representative images from control **(E)** and Stat92E RNAi **(F)** RS tumors induced in larvae with control background. **(G)** Quantification of data presented on E, F. 8-13 eye discs/genotypes were analyzed, n=13 (E), 8 (F). **(H, I)** Representative images from Atg13 mutant **(H)** and Stat92E, Atg13 double deficient **(I)** RS tumors induced in larvae with control background. **(J)** Quantification of data presented on H, I. 5-8 eye discs/genotype were analyzed, n=8 (H), 5 (I). **(K, L)** Representative images from Vps39 mutant **(K)**and Stat92E, Vps39 double deficient **(L)** RS tumors induced in larvae with control background.**(M)** Quantification of data presented on K, L. 8 eye discs/genotype were analyzed, n=8 (K, L).

Tumors can also induce autophagy in host tissues, therefore we were also interested in whether the loss of distinct steps in autophagy may differentially affect JAK/STAT signaling in non-transformed tissues. Hence, we generated RS tumors in larvae where the entire body (including both tumor and host tissues) was mutant for either Atg13 or Syntaxin17 and tested the Stat92E activity in their body wall muscles by western blot. Similarly to its tumor specific effect, systemic loss of Atg13 resulted in decreased levels of Stat92E driven GFP in muscle lysates compared to control or Syntaxin17 mutant ones **(Supplementary Figure S2)**. These findings indicate that in the presence of RS tumors, autophagosome formation, but not autophagosome clearance and autolysosomal degradation, is critical for the maintenance of Stat92E signaling both in tumor and host tissues.

### Stat92E inhibitor Su(var)2-10 is regulated by autophagic sequestration in RS tumors

As JAK/STAT signaling was downregulated in Atg13 but not in Vps39 deficient RS tumors we assumed that autophagosome formation may regulate JAK/STAT signaling by limiting the availability of a JAK/STAT inhibitor in the cytoplasm. Importantly, a recent study in *Drosophila* identified PIAS family protein homolog Su(var)2-10, a known inhibitor of STAT family proteins, as a specific cargo of autophagy^10^. To assay whether inhibition of Stat92E signaling in Atg13 mutant tumors is due to the accumulation of Su(var)2-10, we carried out western blot analysis on day 7 control and Atg13 or Vps39 mutant RS tumors. Importantly, Su(var)2-10 levels were increased in both Atg13 and Vps39 mutants, however this was accompanied with lower Stat92E activity only in Atg13 mutants **(Figure 4A)**. To determine how elevated Su(var)2-10 levels did not lead to a detectable decrease in Stat92E activity, we carried out immunolabeling in effort to determine the subcellular localization of Su(var)2-10 in Vps39 mutant RS tumors that additionally expressed the autophagic vesicle marker mCherry-GFP-Atg8a. We observed that majority of Su(var)2-10 localized to the nucleus, while cytoplasmic Su(var)2-10 frequently showed punctate pattern that colocalized with Atg8a reporter in Vps39 mutant tumors **(Figure 4B)**. Hence, although Su(var)2-10 accumulates both in Atg13 and Atg39 mutants, it is likely that it is already trapped in membrane bound autophagosomes in Vps39 and thereby isolated from the cytoplasm.

**Figure 4.**
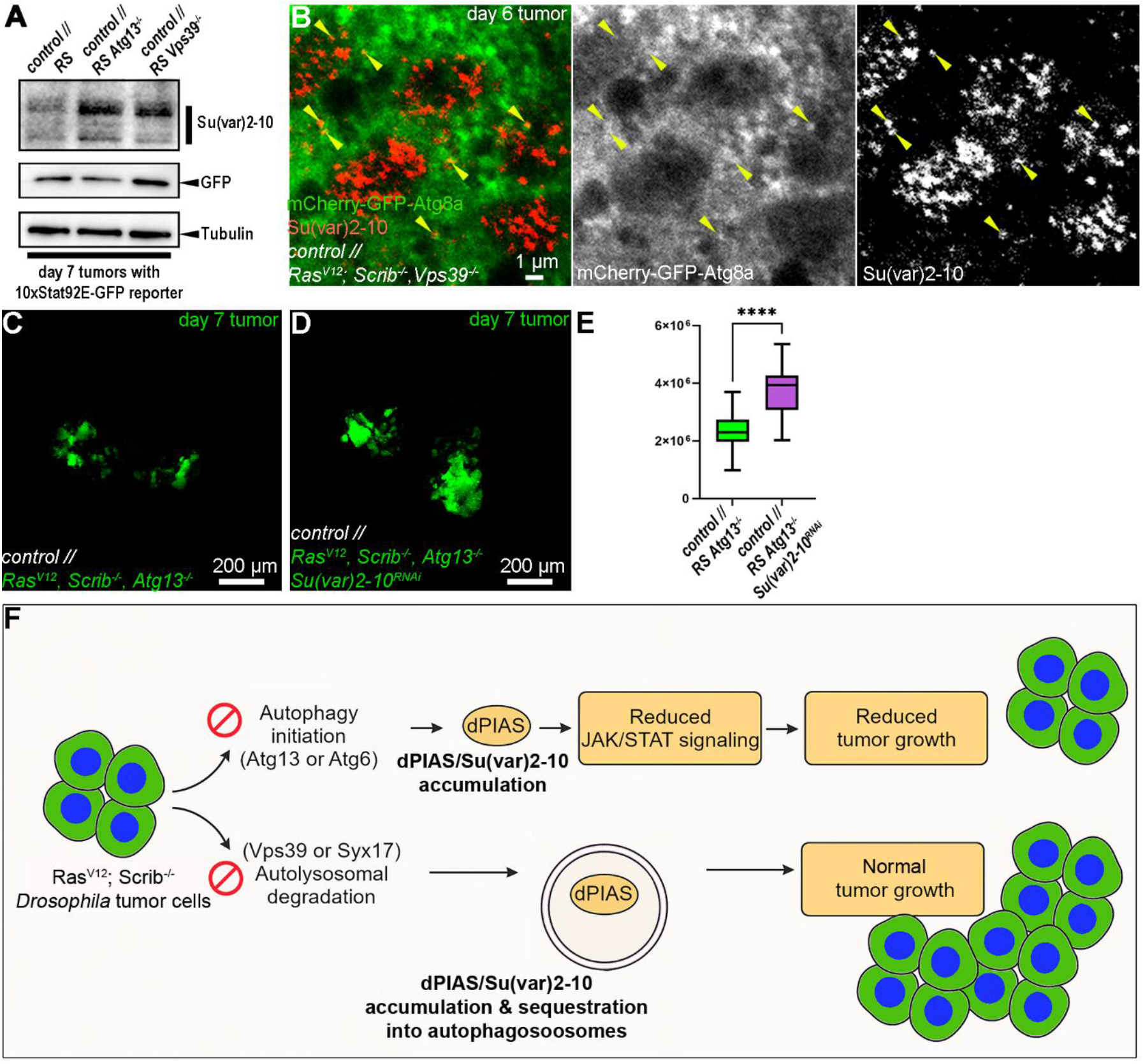
Autophagy regulates Stat92E by sequestrating Su(var)2-10. **(A)** Western blotting of day 7 tumors to detect levels of Stat92E inhibitor Su(var)2-10 in RS Atg13^-/-^ and RS Vps39^-/-^ and control RS tumors. **(B)** Immunolabeling of Su(var)2-10 together with visualization of GFP-Atg8a positive autophagosomes in RS Vps39^-/-^ tumors. Arrowheads as reference points where overlap between channels was observed. **(C, D)** Representative images of Atg13 mutant **(C)** and Su(var)2-10 Atg13 double deficient tumors **(D)** induced in larvae with control background. **(E)**Quantification of data presented on C, D. 14-19 tumors/genotypes were analyzed, n=14 (C), 19(D), unpaired T-test, ****: p<0,0001. **(F)** Our suggested model showing how loss of distinct steps of autophagy affects JAK/STAT signaling in RS tumors. By being an autophagic cargo, Su(var)2-10 (abbreviated as dPIAS here) is normally degraded in autolysosomes, while loss of genes required for early steps of autophagy (Atg13, Atg6) results in the persistence of Su(var)2-10 (abbreviated as dPIAS here) in the cytoplasm, which eventually leads to decreased JAK/STAT signaling and reduced tumor growth. In contrast, upon the loss of autophagosome-lysosome fusion (Vps39 and Syx17 mutant tumors) the persisting dPIAS/Su(var)2-10 is already sequestered into autophagosomes, thereby JAK/STAT signaling and tumor growth remains intact.

To assay whether the accumulation of Su(var)2-10 sufficiently explains the decreased size of Atg13 mutant tumors we knocked down Su(var)2-10 in a tumor specific manner and tested how it affected their growth. Most importantly, loss of Su(var)2-10 partially, but significantly restored the growth of Atg13 mutant tumors **(Figure 4C, D)**. This finding suggests that elimination of Su(var)2-10 from the cytoplasm of cancer cells, by sequestration into autophagosomes – and independently of its subsequent degradation – is critical to maintain tumor progression.

## Discussion

In our study we provided evidence that distinct steps of autophagy are differentially required for tumor progression. By blocking early steps of autophagosome formation (Atg13 and Atg6 mutants) specifically in tumors, or even in the entire body, we observed substantial reduction in the growth of RS tumors. While the lack of autophagosome-lysosome fusion (Vps39 and Syx17 mutants) and hereby autophagic degradation in the tumor tissue or in the whole body did not affect tumor size. Our observation that Atg13, but not Vps39, mutant tumor cells show reduced proliferation and increased apoptosis indicates that RS tumor cells primarily depend on early steps of the autophagic process. Specifically, autophagosome formation appears to be critical, whereas cargo degradation and nutrient release may not serve as the major tumor growth promoting function of autophagy in the RS tumor context.

Our finding that JAK/STAT signaling is downregulated in Atg13 and Atg6 but not Vps39 mutants suggests that autophagy facilitates the activity of this signaling pathway in a degradation independent manner. In parallel we also observed the accumulation of STAT inhibitor Su(var)2-10. This is in line with a recently published study^10^ showing that Su(var)2-10 is a specific cargo of autophagy, and autophagy mediated degradation of Su(var)2-10 is required for upregulation of Stat92E signaling in glial cells upon axon injury. As autophagosomes cannot form in Atg13 mutants it is likely that the accumulated Su(var)2-10 is contributing to the downregulation of Stat92E signaling and thereby the decreased progression of Atg13 deficient RS tumors. In support of this their growth could be partially restored by the concomitant silencing of Su(var)2-10. However, as autophagosomes can still form in Vps39 mutant tumors, it is likely that accumulated Su(var)2-10 is sequestered and trapped in autophagosomes in Vps39 tumors **(Figure 4F)**, a scenario further supported by our finding that punctate Su(var)2-10 signal colocalizes with Atg8a positive autophagosomes in Vps39 mutant tumors. This suggests that sequestration, but not degradation, of the autophagic cargo could be fulfilling the regulatory function of autophagy on Stat92E signaling, highlighting a novel non-degradative role of autophagy in tumor progression.

Tumor induced autophagy in the host tissues of larvae that carry RS tumors does correlate with an increase of amino acids and sugars in the hemolymph, which are likely assimilated into the tumor biomass^6,27^. Earlier observations that the loss of Atg13 or Atg14 in the whole larva substantially blocks tumor growth raised the possibility that this massive mobilization of nutrients from host tissues is mediated by autophagy^8^, and the role of autophagy in releasing nutrients from host tissues was also described in mice^7^. However, one of our most interesting finding was that RS tumors can progress normally in Syx17 mutant larvae, in which autophagosome-lysosome fusion and autophagic degradation was blocked in the whole body, suggesting that the autophagic degradation in host tissues may not be the only process that is directly involved in that nutrient release. Our further finding, that Stat92E signaling is downregulated in the body carcasses of Atg13, but not Syx17 mutant larvae carrying RS tumors, suggests that autophagosome formation may support the activation of JAK/STAT inflammatory signaling in the entire body. Importantly, it has been reported both in *Drosophila* and mammalian in vivo tumor models that elevated JAK/STAT signaling can induce catabolic processes in host tissues^28^. This raises the possibility that autophagy may support nutrient release from host tissues by facilitating JAK/STAT inflammatory signaling.

The interplay of autophagy and JAK/STAT have been reported in a tumor setting multiple times. Autophagy is contributing to the activation of JAK/STAT by promoting IL-6 cytokines in breast^23,29^ and pancreatic ^24^ cancer cells, while in another study showed that inhibition of STAT3 signaling in other cancer cell lines can downregulate autophagy decreasing the expression of autophagy-related gene LC3^30^. The recent discovery that Su(var)2-10 is a cargo of autophagy^10^, opened new perspectives in understanding the interplay of autophagy and Stat92E signaling, however the significance of this phenomenon in a tumor setting was not tested before. PIAS family proteins, the mammalian homologs of Su(var)2-10, are SUMO-ligases and have multifaced role in tumor progression^31^. However, as PIAS proteins have multiple targets in addition to members of JAK/STAT signaling, their negative or positive effect on tumor progression is highly dependent on the types of the PIAS proteins and the tumors. PIAS3 is a well-characterized negative regulator of JAK/STAT and low levels of PIAS3 is associated with elevated JAK/STAT signaling and cancer progression^32-34^, while elevated of PIAS3 results in diminished progression of multiple cancer cell types^35,36^. Our findings that the level of Su(var)2-10 is elevated in autophagy deficient RS tumors, and that cancer cell specific knock down of Su(var)2-10 can partially restore the growth of Atg13 mutant tumors suggests that autophagy mediated sequestration and clearance of Su(var)2-10 is an important link between autophagy and the activation of JAK/STAT signaling in RS. This is further supported by our observation that in contrast to control or Vps39 mutant tumors, whose progression is highly dependent on the presence of Stat92E, silencing of Stat92E does not cause any further decrease in Atg13 mutant tumor size. However, as the Su(var)2-10 and Atg13 double deficient tumors did not gain back their full growth potential, we cannot rule out that there can be other regulatory mechanisms between autophagy and tumor growth, likely through acting on other signaling processes and that the lack of autophagy mediated nutrient release may also contribute here to some extent.

Overall, our research demonstrated that the early steps of autophagosome formation and cargo sequestration, but not autophagosome-lysosome fusion and the subsequent cargo degradation, are critical for the growth of RS tumors and the tumor-associated JAK/STAT inflammatory signaling, both in the tumor and the host tissues. This highlights that the autophagic degradation-mediated nutrient release is not essential for tumor growth. Moreover, our findings point out that the characterization of alternative functions of autophagy, such as the sequestration of critical signaling molecules, that are not directly linked to autolysosomal degradation, can bring us closer to a fuller understanding of the role of autophagy in tumor progression.

## Materials and Methods

### Fly stocks and genetics

*Drosophila* stocks were maintained on standard fly food containing agar, corn flour, sugar, CaCl_2_, yeast, and antifungal agents nipagin and 2-Phenylphenol dissolved in ethanol. Flies were kept at a constant 25 °C. Fly stocks were obtained from Bloomington *Drosophila* Stock Centre (Indiana, US) except the following mutant alleles: Atg13 ^Δ81^ (from Thomas Neufeld), Atg6 ^1^ (from Eric Baehrecke), Vps39^Δ1^ (from Ole Kjaerulff), *Syx17*^*LL06330*^ (from Kyoto *Drosophila* Stock Center, Kyoto, Japan). Detailed Genotypes of larvae used in experiments are listed in **Supplementary Table S1**.

Tumorigenesis in the larval eye discs was induced with MARCM (Mosaic Analysis with Repressible Cell Marker) technique. Stocks containing *UAS-Ras*^*V12*^ oncogene along with a mutant allele (*Scrib*^*1*^) for *Scribble* cell polarity gene were crossed with ey-FLP containing MARCM lines (see detailed genotypes **Supplementary Table S1**). Crossings were transferred to new tube every 24 hours, to keep track of age of flies. According to the type of experiment the larvae were of age 6 or 7 days.

As tumors were induced in larvae with various genetic background (control, Atg13 or Syx17 mutant) the genotypes in each experimental panels were indicated by the following formula: *genotype of the host tissues // genotype of the tumor*.

### Tumor volume analysis

Eye disc tumors were dissected from larvae in PBS and fixed in 4% PFA dissolved in PBS for 20 minutes. Samples were washed in PBS three times for 5 minutes. Finally, samples were mounted on a glass slide in glycerol. 0,22 mm spacers were used between the slide and the coverslip to avoid the compression of tumors. Samples were imaged by carrying out optical sectioning (Z-stack, interval: 3 µm, 10x magnification) with a Zeiss AxioImager M2 fluorescent microscope, equipped by Apotome2 confocal unit (Zeiss) and Orca-Flash 4.0 LT3 digital sCMOS camera (Hamamatsu Photonics).

### Immunohistochemistry and fluorescent reporters

Larval eye disc tumors were dissected in PBS and fixed with 4% PFA-PBS for 20 minutes. This was followed by 3 x 5 min washing with 0.5% Triton-X-100-PBS (PBTX). Primary antibodies were diluted in 5% FBS-PBTX and incubated overnight at 4°C. The antibodies were washed out by 3 x 5 min washing in PBTX. Then, samples were incubated in secondary antibody solution (secondary antibodies diluted 5% FBS-PBTX) for 3 hours at room temperature. Antibodies were washed by 2 x 5 min PBTX and 2 x 5 min PBS washing steps. Then samples were mounted in glycerol and analyzed with Zeiss Axioimager M2 fluorescence microscope on 20x (anti-Dcp1, anti-PH3) and 63x (anti-Atg8a) magnification. Colocalization of mCherry-GFP-Atg8a reporter and anti-Su(var)2-10 immunostaining was analyzed with Olympus FV3000 Laser Scanning Confocal Microscope, on 40x magnification. The following primary antibodies were used: rat anti-Atg8a (1:500)^20^; rabbit anti-P-H3 (Millipore, #06-570) (1:300); rabbit anti-cleaved-Dcp-1 (Cell Signaling Technologies, 95785) (1:200); chicken anti-GFP (Invitrogen, #A10262); mouse anti-Su(var)2-10 (from Julius Brennecke, IMBA, Vienna, Austria) (1:100)^37^. The following secondary antibodies were used: anti-Chicken DyLight™ 488 (Invitrogen, #SA5-10070); anti-Rabbit Alexa Fluor™ 568 (Life Technologies; # A11036); anti-Mouse Alexa Fluor™ 647 (Invitrogen; #A21236).

For studying STAT92E activity day 6 tumors were dissected from larvae carrying STAT92E driven GFP reporter and were fixed in 4% PFA-PBS for 20 minutes. This was followed by 2x rinse and 1x 15 minutes wash in PBS. Then, samples were mounted and imaged in the same way as it was described at immunostaining.

### Western blot

For western blot experiments larval eye disc tumors or larval muscle tissue were dissected in PBS and collected in ice cold lysis buffer (50 mM Tris-HCL pH 8.0, 50 mM KCl, 10 mM EDTA, 1% Triton X-100) supplemented with protease inhibitors (PMSF, Aprotinin, Leupeptin) and phosphatase inhibitors (NaF, Na_3_VO_4_). Samples were homogenized with micro pestles and spun on microcentrifuge (13 000 rpm). Total protein concentrations were measured by using Bradford reagent (5000006, Bio-Rad). Laemmli buffer was added to the remainder of sample and heated on 95 °C for 5 minutes in a thermoblock. Finally, the samples were centrifuged for 2 minutes on 13 000 rpm and stored at -20 °C until further use.

Western blot experiments were carried out by following standard protocols. 5 µg of each sample were loaded on polyacrylamide gels and SDS-PAGE was carried out with Mini-PROTEAN® Tetra Vertical Electrophoresis Cell (1658004, Bio-Rad). Proteins from the gels were transferred to 0,45 µm LF PVDF membranes using transfer kit (1704275, Bio-Rad) with Trans-blot® Turbo™ system (1704150, Bio-Rad). Membranes were incubated in 0,5 % casein blocking solution for one hour at room temperature, followed by overnight incubation with primary antibodies diluted in 1:1 mixture of casein : TBST (0,1% Tween -TBS) at 4°C. Membranes were washed three times in TBST for 10 mins with subsequent incubation with secondary antibodies diluted in 1:1 casein : TBST for one hour at room temperature. Secondary antibodies were washed out by three 10 minutes washing steps in TBST. Finally, the membranes were developed with HRP substrate solution (WBKLS0500, Millipore). Bands were detected with ChemiDoc™ Imaging System (12003153, Bio-Rad). The following primary antibodies were used: rat anti-Atg8a (1:5000)^20^; rat-anti-mCherry (1:5000)^18^; rat anti-GFP (1:4000)^38^; rabbit anti-ref(2)p (1:2000); mouse anti-Su(var)2-10 (from Julius Brennecke, IMBA, Vienna, Austria) (1:1000) ^37^; mouse anti-Tubulin (DSHB, #AA4.3) (1:2000). The following secondary antibodies were used: anti-Rat HRP (Sigma-Aldrich, #A9037) (1:4000); anti-Mouse HRP (Sigma-Aldrich, #A9044) (1:10000); anti-Rabbit HRP (Millipore, #AP187P) (1:2500

### Image processing and statistics

Image analyses were made by using ImageJ software and data was evaluated with Graphpad Prism 10. The applied statistical probes are described in their corresponding figure. Normality of data distribution was assessed using the Shapiro-Wilk test. Unpaired T-tests were carried out in case of side-by-side comparison for two datasets with normal distribution. One-way ANOVA or Kruskal-Wallis tests were used for multiple comparisons, when all datasets had or did not have normal distributions respectively.

For tumor volume analysis, Z-stack images were processed with a custom macro in ImageJ. First, a 3D Gaussian blur with a sigma of 2 voxels was applied to the green channel representing the tumor across the whole stack. Next, Otsu thresholding was performed with the “dark background” option enabled, and the resulting images were converted into a binary mask. Tumor volume was then measured using the 3D Objects Counter, where only objects ranging from 20 to 100,000,000 voxels were considered, and the fixed threshold intensity was set to 128.

Z-stack (with intervals 1,25 µm) images of PH3 immunohistochemistry experiments were acquired, and for each sample a single image was selected for further analysis. The selection was made by choosing the third image of the Z-stacked image inward from the tissue surface toward the deeper tissue layers. A custom macro was then created in ImageJ to process the images. To preprocess the images the channels were split, after which a Gaussian blur was applied to the red channel containing the PH3 signal that marks dividing cell nuclei. We used the moments automatic thresholding to convert into a binary mask, and refinement with watershed segmentation to separate all the cell nuclei even if they have merged in the image. Nuclear puncta were then detected in the red channel, and their centroids (X,Y) were extracted. The green channel, corresponding to the tumor area, was subsequently processed by applying a Gaussian blur, followed by Otsu auto thresholding and conversion to a binary mask. The GFP-positive area was quantified by particle analysis, and the number of PH3 puncta located within the tumor area was determined. Finally, the percentage of PH3 puncta within the tumor area was calculated for each image of each genotype.

For the analysis of apoptotic cells within tumor areas, Z-stack images were processed using a custom ImageJ macro. The channels were split into separate images corresponding to the DCP1 apoptosis marker (Red channel), and tumor marker (Green channel).

To quantify tumor regions, the green channel was first duplicated and processed with a 3D Gaussian blur. Each slice of the stack was thresholded manually using the ImageJ thresholding tool, and the resulting images were converted into binary masks. The same procedure was applied independently to the red channel.

For each slice, the green binary mask was used to define the tumor area as a region of interest (ROI). These ROIs were applied to the corresponding slices of the red binary mask, and apoptotic signals were quantified within the tumor regions using the Analyze Particles function with a size range of 10–2,000,000 pixels^2^. This allowed the calculation of apoptotic particles located within tumor areas.

For Atg8a puncta measurement we used a custom macro. Z-stack images were imported as hyperstacks, and individual channels were separated.

The GFP channel was converted to 8 bit and denoised with Gaussian blur (σ = 5) applied across the stack and thresholded manually on a representative middle slice (5th slice from 9 slices). ROIs were generated for the middle three slices (slices 4, 5 and 6).

Far-red channel images were preprocessed with Gaussian blur. Thresholding was done manually on the middle slice (5th slice). Previously defined ROIs were used on their respective far-red slice, and far-red area fraction within each GFP ROI was measured.

## Funding & Acknowledgements

This research was funded by the National Research, Development and Innovation Office of Hungary (OTKA FK_142508 to ST and EKÖP-25-2-I-ELTE-80 to NN), the Hungarian Academy of Sciences (BO/00400/23 to ST), and the Excellence Fund of Eötvös Loránd University (EKA_2022/045-P302-1 to ST). FOF was supported by a Norwegian Research Council Grant “Decoding tumor cell invasive switching” Project No.: 324447.

We thank Ivett Répássy for her technical assistance.

## Author contributions

AR, NN, FOF and ST designed and provided the resources for the research. AR, NN, DK and BSB performed the experiments and analyzed the data. BSB wrote scripts and algorithms in Python and ImageJ. AR, FOF and ST wrote the manuscript.

## Conflict of interest

The authors declare that no conflict of interest exists.

## Notes

### Competing Interest Statement

The authors have declared no competing interest.

