## Supplementary Figures S1-2 for "Autophagy promotes tumor growth through facilitating JAK/STAT signaling in a lysosomal degradation independent manner"

### Supplementary Material

#### Supplementary Figures and Supplementary Figure Legends

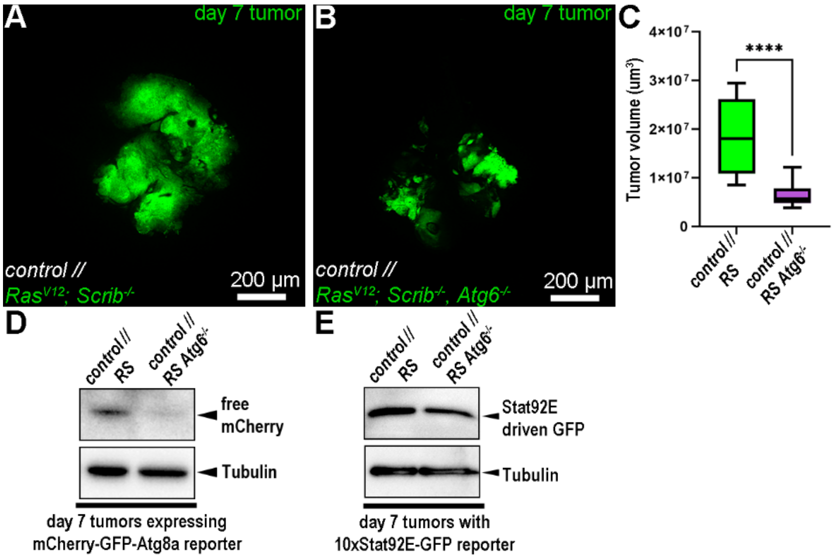


**Supplementary figure S1. Loss of Atg6 results in decreased progression and JAK/STAT signaling. (A, B)** Representative images about control **(A)** and Atg6 mutant **(B)** RS tumors that were induced in larvae with control background. **(C)** Quantification of data presented on A, B. 12 tumors/genotype were analyzed, n=12 (A, B), unpaired T-test, ****: p<0,0001. **(D)** Western blotting of day 7 tumors reveals that free mCherry, the lysosomal cleavage product of mCherry-GFP-Atg8a reporter, is not generated both in Atg6 deficient tumors. **(E)** Western blotting of the lysates of day 7 tumors reveals that Atg6 deficient tumors show reduced amount of Stat92E activity driven GFP.

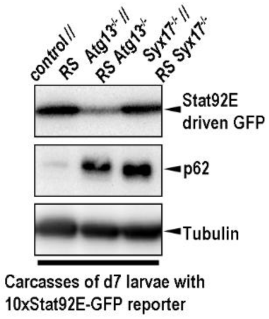


**Supplementary Figure S2. JAK/STAT signaling is downregulated in the muscles of Atg13 mutant larvae that carry RS tumors.** Western blotting on the lysates of carcasses from day 7 larvae carrying RS tumors to compare the effect of loss of Atg13, but not Syx17 in the whole body on Stat92E activity in the body wall muscles. Selective autophagy target protein p62 was used to monitor autophagic degradational capability of control and mutant genotypes.

**Supplementary Table S1. Detailed genotypes in each Figures.**
